# Surviving under Ice: Insights into gene expression changes during ice encasement in timothy (*Phleum pratense* L.)

**DOI:** 10.1101/2024.04.11.589102

**Authors:** Akhil Reddy Pashapu, Sigridur Dalmannsdottir, Marit Jørgensen, Odd Arne Rognli, Mallikarjuna Rao Kovi

## Abstract

The predicted increase in frequency and duration of winter warming episodes (WWEs) at the higher northern latitudes is expected to negatively impact the forage production in this region. The formation of non-permeable ice cover due to WWEs creates hypoxic or anoxic conditions for plants leading to severe winter damages. Knowledge about molecular mechanisms underlying various winter stresses is crucial to develop cultivars with better winter survival under changing climatic conditions. In the current study, we aimed at identifying genes involved in ice encasement stress responses by RNAseq in the perennial forage grass timothy (*Phleum pratense* L.) and study gene expression differentiation due to field survival using cultivars with diverse genetic backgrounds. The LD_50_ estimates varied across cultivars and material. The expression of ethylene-responsive transcription factors, alcohol dehydrogenase, pyruvate decarboxylase, sucrose synthase, dehydrins, and heat shock proteins were highly upregulated under ice encasement conditions. Functional analysis of differentially expressed genes revealed that the upregulated genes were involved in glycolysis, pyruvate metabolism, carbon metabolism, and biosynthesis of amino acids while genes involved in photosynthesis, phenylpropanoid biosynthesis and flavonoid biosynthesis pathways were downregulated. The results from the current study indicate a substantial overlap of ice encasement stress responses with those of hypoxia and freezing stresses. In addition, the potential strategies leading to higher ice encasement tolerance of timothy are outlined. Furthermore, differences in gene expression between field survivors and the original plant material, and differences between ice encasement responses of northern-adapted and southern-adapted cultivars are briefly discussed.

## Introduction

The increase in global temperatures due to climate change is projected to create variable weather patterns across the globe (IPCC, 2014, 2022), affecting both agriculture and natural ecosystems. In particular, at higher latitudes, the increase in temperature is predicted to be greater during winter than in summer (Overland et al., 2011; IPCC, 2014). Though the climate change at higher latitudes is projected to be beneficial in the long run due to the crop shifts northward (Olesen et al., 2007; Franke et al., 2022), the concomitant alterations in winter weather are likely to have a negative impact on forage production in this region (Rapacz et al., 2014; Bjerke et al., 2015). For instance, Dalmannsdottir et al. (2017) and Jørgensen et al. (2020) reported that an increase in autumn temperature and precipitation might impair cold acclimation in grasses, thus reducing their capacity to tolerate frost. Furthermore, reports indicate that there was an increase in the frequency and duration of winter warming episodes (WWE) over the past few decades, a trend which is expected to amplify in the future (Thorsen and Höglind, 2010; Johansson et al., 2011; Vikhamar-Schuler et al., 2016). Snowmelt from these WWEs could expose the plants to ambient air temperatures, increasing the risk of winter kill. In addition, WWEs can also lead to the formation of nonpermeable ice cover (ice encasement) causing hypoxic or anoxic conditions for plants. Ice encasement for an extended period can be detrimental to the survival of plants (Gudleifsson, 2009), as a result, tolerance to ice encasement/ice cover has become critical for the winter survival of perennial grasses. Moreover, the WWEs could subject the plants to winter desiccation due to frozen ground (Bjerke et al., 2014, 2015), and may also pose the risk of premature dehardening (Inouye, 2008). Not surprisingly, there have been several reports from farmers about increased winter kill and reduced persistence of perennial grasses at higher latitudes in recent years (Jørgensen, 2017). This highlights the urgent need to develop new cultivars with better tolerance to various stresses under novel winter conditions, to ensure stable and sustainable forage production in this region.

Stress responses in plants are complex, involving sensing, signalling and subsequent transcriptional and translational alterations (Lamers et al., 2020; Zhang et al., 2022). Hence, knowledge about genes, gene regulation and biochemical pathways involved in tolerance to various winter stresses is crucial to breed new cultivars with better winter survival. Over the years, several transcriptome studies aimed at understanding the molecular stress responses in plants have revealed key insights into genes and pathways involved in freezing tolerance (Fu et al., 2016; Zuther et al., 2018; Xu et al., 2019; Wu et al., 2022c), snow mould resistance (Kovi et al., 2016; Tsers et al., 2021) and desiccation tolerance (Verdier et al., 2013; Chávez Montes et al., 2022). However, to date, no such study has been carried out to dissect the molecular mechanisms underlying ice encasement tolerance in plants, emphasizing the need to address this knowledge gap.

Ice encasement tolerance (ICET) can be defined as the ability of plants to withstand ice cover for an extended period without incurring irrecoverable damage. ICET is expressed in terms of LD_50_, i.e., the number of days under ice cover that kill 50% of the plant population (Gudleifsson and Bjarnadottir, 2014). Despite decreased photosynthetic activity (Höglind et al., 2011; Borawska- Jarmułowicz et al., 2014) and metabolic turnover (Gudleifsson, 2009) under freezing conditions, perennial plants respire albeit at reduced rates (Ögren, 2000) throughout the winter to fuel the cellular processes which protect the cells from stress or helps them mitigating it. Thick non-permeable ice cover (ice encasement) may develop at the end of WWEs restricting the gaseous exchange between the plant and the surrounding environment, leading to hypoxic or near-anoxic conditions under which plants are eventually forced to switch to anaerobic respiration (fermentative pathway) (Drew, 1997; Höglind et al., 2010; Licausi et al., 2011; Gudleifsson and Bjarnadottir, 2014). The anaerobic fermentative pathway is energy inefficient; yielding 2 mols of ATP for each mol of glucose compared to 32 mols of ATP under aerobic respiration (Vartapetian and Jackson, 1997), and leads to the production of by-products like CO_2_, ethanol, malate and lactate etc. (Gudleifsson, 1994, 1997) whose accumulation at higher levels can be phytotoxic (Andrews and Pomeroy, 1979).

Plants under ice casement conditions face a unique set of challenges. The initial set of challenges; freezing stress, dehydrative stress, mechanical stress and hypoxic stress originate due to the formation of the thick non-permeable ice cover, while the later set of challenges; starvation due to depleted carbohydrate reserves, phytotoxicity due to accumulation of anaerobic byproducts and snow mould infestation arise because of prolonged ice cover. Among them, freezing stress and hypoxic stress can be considered the primary stressors, while the remaining are secondary effects resulting from these primary stressors. Thus, it can be hypothesized that molecular responses under ice encasement overlap with those under freezing stress and hypoxic stress. During freezing stress, plants have to cope with the formation of ice in the apoplast, dehydrative stress, protect cell membranes from oxidative damage and synthesize proteins which stabilize cellular functions (Thomashow, 1998; Parvanova et al., 2004; Sandve et al., 2011; Ritonga and Chen, 2020), whereas under hypoxic stress, which usually occurs due to flooding/submergence, mitigating the energy imbalances due to limited ATP synthesis by oxidative phosphorylation, and efficient management of energy reserves while preventing the accumulation of anaerobic byproducts to toxic levels are crucial (Igamberdiev and Hill, 2009; Bailey-Serres and Voesenek, 2010; Loreti et al., 2018). Given the nature of the stress, the molecular mechanisms underlying ice encasement tolerance must be tightly regulated to strike a fine balance between various stress responses and are therefore complex.

Many aspects of submergence/flooding and ice encasement stress are similar in plants (Gudleifsson, 1993). Thus parallels can be drawn between their molecular stress responses. However, ice encasement poses an additional set of challenges (freezing conditions, accumulation of metabolites) and is therefore a more severe form of stress compared to submergence/flooding. The main differences between ice encasement and submergence are i) under ice encasement oxygen deprivation is accompanied by freezing stress (< 0°C), therefore metabolic rates are highly depressed compared to flooding/submergence, ii) the high impermeability of ice to gases (Hemmingsen, 1959) leads to faster development of anoxia, and iii) unlike during flooding/submergence, ice prevents continuous leaching of anaerobic by- products, and this might lead to their accumulation to toxic levels (Andrews, 1977; Andrews and Pomeroy, 1977, 1979; Gudleifsson, 1997). It is, therefore, reasonable to postulate that plant species which are frequently exposed to ice encasement have likely evolved an additional repertoire of molecular mechanisms to mitigate the unique stressful conditions under ice while exhibiting substantial similarities with flooding/submergence stress responses.

Apart from hypothesizing the similarities with submergence/flooding, the molecular mechanisms underlying ICET largely remain unknown to date. Several studies on winter cereals and forage grasses with different ice encasement tolerances have provided valuable insights into the physiological responses of plants to ice encasement (Andrews and Pomeroy, 1977, 1979; McKersie et al., 1982; Gudleifsson, 1994, 2013; Höglind et al., 2010). These studies revealed huge variation in ICET within and between species (Gudleifsson et al., 1986; Gudleifsson, 2010; Höglind et al., 2010). In addition, they also indicated that the accumulation of anaerobic byproducts to toxic levels is the primary reason for cell death under ice encasement conditions rather than oxygen deficiency, freezing temperatures, dehydration, mechanical stress from ice or depletion of carbohydrate reserves (Rakitina, 1970; Andrews and Pomeroy, 1977; Pomeroy and Andrews, 1978; McKersie et al., 1982). Based on these findings, prevention of anaerobic byproduct accumulation to toxic levels and (or) mitigating their effect on cellular processes could be the key factors determining survival under prolonged ice encasement in overwintering grasses. On the contrary, only a few studies have delved into changes at the molecular level under ice encasement conditions. A study from Andrews (1997) probed the changes in the enzyme activity of ATP-dependent 6-phosphofructokinase (PFK), PPI-dependent phosphofructotransferase (PFP), pyruvate kinase (PK), pyruvate decarboxylase (PDC) and alcohol dehydrogenase (ADH) in winter wheat (*Triticum aestivum*), timothy (*Phleum pratense* L.) and Bering hairgrass (*Deschampsia beringensis*) under ice encasement conditions. In addition, a couple of studies examined the changes in the expression of genes encoding glyceraldehyde-3-phosphate dehydrogenase (GAPDH), ADH and lactate dehydrogenase (LDH) under low-temperature oxygen-deficient conditions in four perennial forage species, alfalfa (*Medicago sativa* L.), red clover (*Trifolium pratense* L.), orchard grass (*Dactylis glomerata* L.), and timothy (Bertrand et al., 2001; Bertrand, 2003). To the best of our knowledge, these are the only studies that investigated changes at the molecular level under ice encasement (or) comparable conditions.

Timothy (*Phleum pratense* L.), a cool-season, hexaploid perennial grass species is the most important forage grass in Northern Norway, a region that is expected to have frequent winter warming episodes due to climate change (Vikhamar-Schuler et al., 2016). The results from previous studies indicate that timothy is among the most ice encasement tolerant forage grass species with significant variation in ICET between different cultivars (Andrews and Gudleifsson, 1983; Gudleifsson, 1994; Bertrand et al., 2001; Höglind et al., 2010). In the current study, timothy cultivars from diverse adaptation backgrounds and surviving plants from field trials under different climatic conditions were used to investigate transcriptome-wide changes under long-term ice encasement conditions. The main objectives of the study were to i) identify genes involved in ice encasement tolerance of timothy; ii) study the effect of selection on ice encasement responses and gene expression iii) identify differences in ice encasement responses between northern and southern adapted timothy cultivars.

## Material and methods

### Plant material and field trials

The plant material used in the current study is the same as described in Pashapu et al. (2024). To briefly summarize, 4-year-old surviving plants of the timothy cultivars Engmo, Noreng, Grindstad, and Snorri were collected from field plots in Tromsø (69.4° N, 19.4° E) and Vesterålen (68.7° N, 15.4° E) in late August/early September 2020. These survivors were tested for ice encasement tolerance along with young plants raised from the original seed lots used to establish the field trials (Figure S1).

### Ice encasement tests, estimation of LD50, and sequencing

Initially, the plants raised from the original seed lots were sown on 27 July 2020 and grown in the greenhouse for 14 days at 18°C, they were later transplanted to tree nursery trays and grown for another 5 weeks at 18°C. The 4-year-old timothy survivors were also planted into nursery trays. Cold acclimation treatment for all plants started on 18th September 2020. The plants were first cold-acclimated under ambient light conditions for 11 days at 9°C, followed by 10 days at 6°C, and finally three weeks at 2°C under 12h light (55 PPFD). The ice-encasement tests were carried out as described in Höglind et al. (2010). The cold-acclimated plants, trimmed to ca. 3 cm top and 1 cm root, were placed into polyethylene boxes and covered with crushed ice before being filled with cold water (3-4°C) and covered with a lid (Figure 1). The boxes were placed in a freezing chamber at −2°C in complete darkness and boxes with the plants were sampled after 24, 38, 52, and 66 days. For genetic studies, plants were placed in separate boxes with two replicates of two seedlings for each entry, which were flash-frozen in liquid nitrogen for subsequent RNA extraction. Another set of boxes with seedlings was used to test the ice encasement tolerance and regrowth, two replicates of eight plants each were used per entry at a given ice encasement stress duration. They were concurrently sampled, thawed at 2°C, transplanted into fertilized peat and scored for survival after three weeks of growth at 18°C under long days conditions (18h). LD_50_ was estimated using glm followed by *dose.p* function from the R package MASS (v7.3.55) (Venables et al., 2002). Cold-acclimated (CA) plants were sampled and used as controls while samples from treatments below LD_50_ were pooled into Treatment 1 (T1) and Treatment 2 (T2). For example, if the LD_50_ of a cultivar is 63, the plant material sampled at 24 and 38 days are pooled together as T1 while material from 52 days was used as T2. RNA was extracted with the Qiagen RNeasy Plant mini kit according to the manufacturer’s guidelines. On-column DNA digestion was performed to remove genomic DNA using Qiagen RNase-Free DNase Set. The RNA yield, purity and integrity were determined by Nanodrop 1100 (ThermoScientific) and Agilent Bioanalyzer 2100 (Agilent Technologies) respectively. The cDNA library preparation and sequencing were outsourced to Novogene Co Ltd (Cambridge, UK) where all samples were paired-end (PE) sequenced using Illumina HiSeq PE150 with an average of ∼ 20 million raw reads per sample.

**Figure 1:**
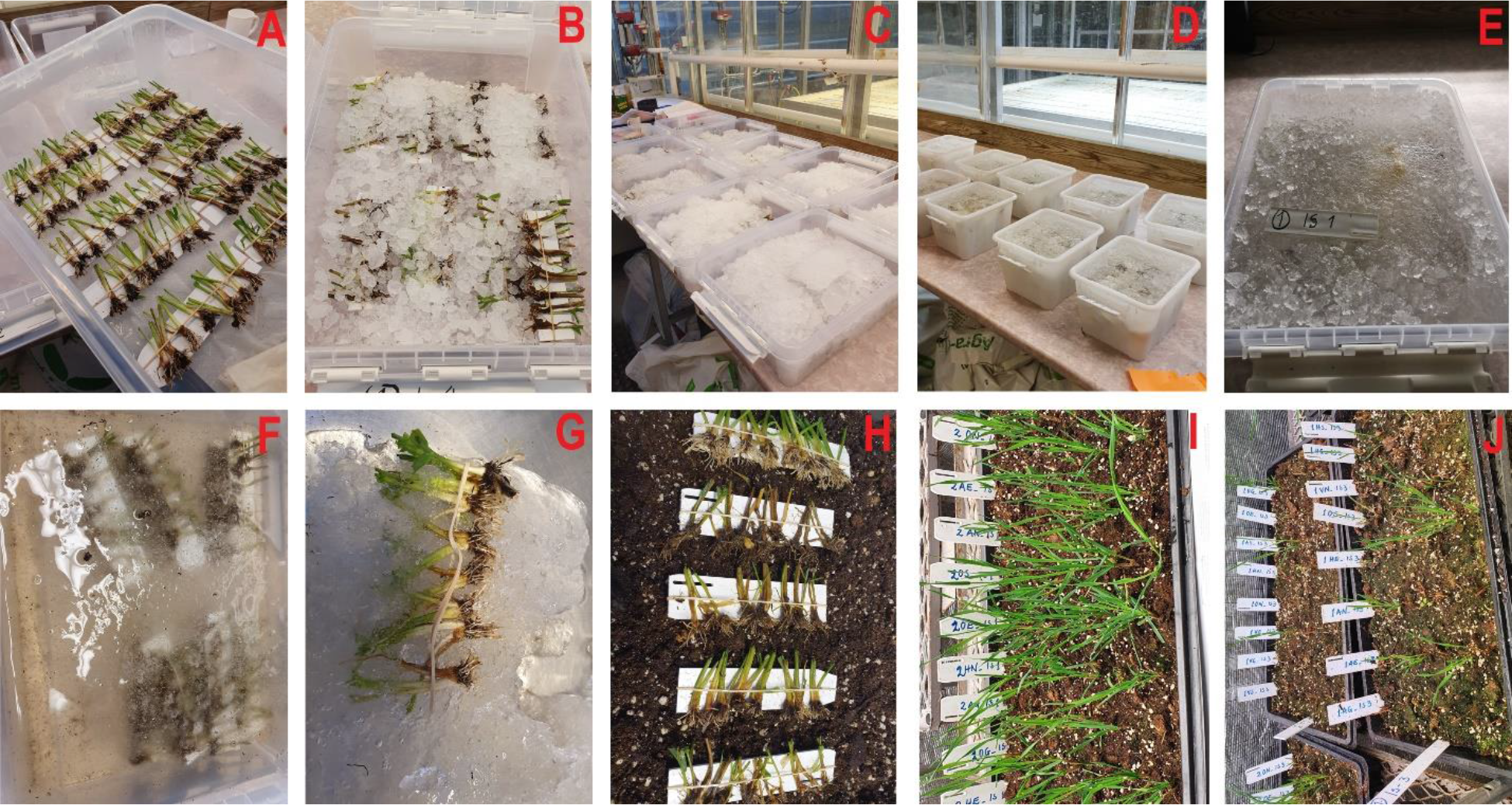
Overview of ice encasement tests. A) Batches of cold acclimated plants. B, C and D) Boxes filled with ice and cold water. E) Boxes ready for transferring to dark cold room. F) Plants encased in ice during the test. G) Sampling after the tests H) Thawing plant material after the tests. I) Regrowth of plants sampled after 24 days J) Regrowth of plants sampled after 52 days.

### Transcriptome assembly, abundance estimation, and functional annotation

After examining the quality of the raw reads with FastQC (v0.11.9) (Andrews, S, 2010) the low-quality bases were trimmed using fastp (0.23.2) (Chen et al., 2018) with options *length_required* 80, *cut_window_size* 4 and *cut_mean_quality* 25. Ribosomal RNA reads were filtered out using SortMeRNA (4.3.6) (Kopylova et al., 2012) against rRNA databases downloaded from https://github.com/sortmerna/sortmerna with an e-value threshold of 1e-5. The cleaned reads were then aligned to a draft reference genome of timothy (unpublished) with STAR (2.7.10b) (Dobin et al., 2013). Initially, reference-guided transcriptomes were assembled for all samples using StringTie (2.2.1) (Kovaka et al., 2019), which were then merged using stringtie merge with options *min_cov* 10 and *min_tpm* 1. The assembly generated by stringtie merge was used as the final reference-guided transcriptome assembly. The transcript sequences were extracted from the reference genome using *rsem-extract-reference-transcripts* script (RSEM) by providing the GTF file generated by stringtie merge. The completeness of the transcriptome was accessed by BUSCO (Benchmarking Universal Single-Copy Orthologs) analysis (5.4.3) (Simão et al., 2015) against the Poales lineage (poales_odb10) and abundance was estimated using RSEM (1.3.3) (Li and Dewey, 2011) with bowtie2 (2.5.1) (Langmead and Salzberg, 2012) as the aligner. The resulting gene level count matrix was used for further downstream analysis. To assign functional annotations based on homology, the transcript sequences were blasted against protein sequences of *Triticum aestivum* downloaded from Ensembl Plants, UniProt, and NCBI using Diamond (2.0.15) (Buchfink et al., 2021) with option *--ultra-sensitive* and e-value cutoff of 1e-5. The GO ids of the annotated sequences were retrieved using the biomaRt R package (v2.50.3) (Durinck et al., 2009).

### Differential expression and functional enrichment analyses

The differential expression and all further downstream analyses were carried out in R (4.2.3) (R Core Team, 2022). The function *filterByExpr* from the edgeR package (Robinson et al., 2010) was used to filter out transcripts with low expression. To validate the experimental setup and identify the main sources of variation, principal component analysis (PCA) was performed based on one thousand highly variable genes. Differentially expressed genes (DEGs) for all contrasts of interest were identified using the DESeq2 (1.38.3) (Love et al., 2014) using the linear regression model ∼0+*MC+MC:Treatment*, where *MC* is the combined factor of material and cultivar. DEGs at group levels were extracted with *results* function by providing the necessary contrast list. For a gene to be considered as differentially expressed it should pass the *padj* < 0.05 and *lfcThreshold* ≥ log2(1.2) filters. Gene Ontology (GO) and Kyoto Encyclopedia of Genes and Genomes (KEGG) pathway enrichment analyses were performed on the sets of upregulated and downregulated DEGs using the topGO (2.50) (algorithm=“lea”, statistic=“fisher”) (Adrian Alexa, 2021) and clusterProfiler (4.6.2) (enrichKEGG function, organism=Taes) (Wu et al., 2021) packages respectively. Enriched GO terms are further reduced based on similarity using the rrvgo package (1.10) (method= “Lin”, threshold=0.7) (Sayols, 2023). All transcripts with hit against Ensembl and NCBI wheat databases are used as background for functional enrichment analyses. DEG expression patterns were visualized by *Heatmap* function from the ComplexHeatmap package (Gu, 2022).

## Results

### Survival trends after Ice encasement tests and LD_50_

The LD_50_ values of timothy cultivars in this study ranged from 37 days to 62 days (Table 1). The plants raised from the original seedlots had higher LD_50_ than their respective field-surviving counterparts across all cultivars. The order of ice encasement tolerance in cultivars across the material was Snorri ≥ Engmo > Noreng > Grindstad. In general, the northern-adapted cultivar Engmo (landrace) had better ICET than the southern-adapted cultivar Grindstad (landrace). The synthetic cultivar Noreng, which is based on genotypes selected within Engmo and Grindstad, had an intermediate ICET compared to its parental cultivars. The cultivar Grindstad had remarkably stable LD_50_ across the material, while substantial declines were observed in the field survivors of the other three cultivars compared to their respective original material (Table 1).

**Table 1:** LD50 of cultivars from different materials. Superscript * indicates skewed LD50 estimates due to missing data.

|  | Engmo | Noreng | Grindstad | Snorri |
| --- | --- | --- | --- | --- |
| **Original seed** | 59.1 ± 2.8 | 47.8 ± 2.6 | 39.6 ± 2.6 | 63.3 ± 2.9 |
| **Vesterålen** | 46.3 ± 2.7 | 29.3* ± 5.2 | 36.8 ± 2.6 | 36.2* ± 3.1 |
| **Tromsø** | 46.9 ± 2.7 | 40.9 ± 2.7 | 36.8 ± 2.7 | 46.9 ± 2.6 |

There were not enough surviving plants in Vesterålen to do a full series of testing. Principal component analysis was performed based on one thousand highly variable genes. PC1 explaining 63% of the variation separating the cold-acclimated (CA) samples and the ice-encased (T1 & T2) samples, while PC2 explaining only 3% of the variation separating the cold-acclimated samples of original and field surviving material (Tromsø & Vesterålen) with no clear separation of ice encased (T1 & T2) samples across different material (Figure 2).

**Figure 2:**
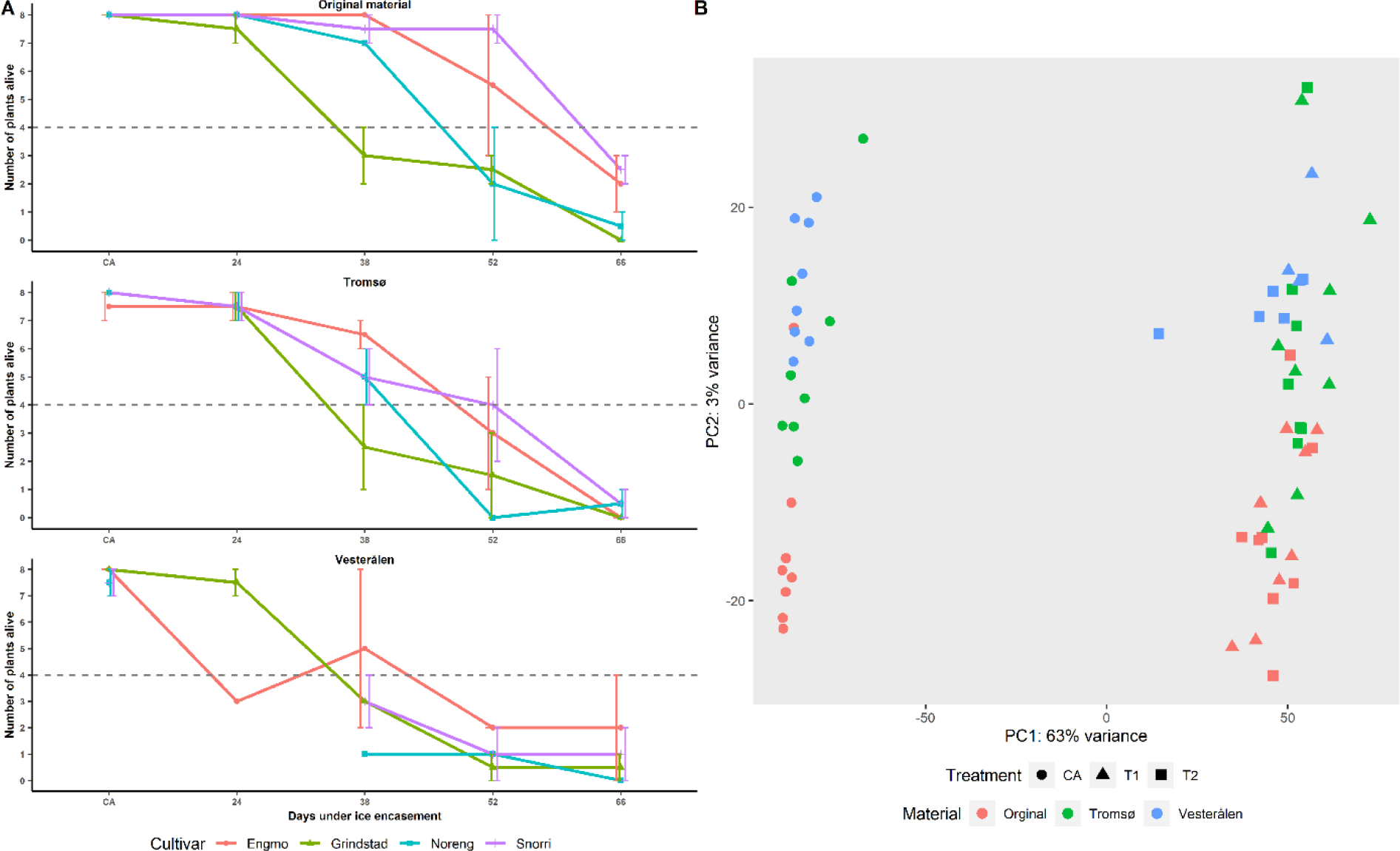
A) Survival graphs after ice encasement tests. Bars denote standard error. X-axis and y-axis denote days under ice encasement and number of plants alive, respectively. B) PCA plot based on one thousand highly variable genes.

### Transcriptome assembly and functional annotation

Trimming low-quality bases and filtering rRNA reads discarded ∼ 3-7% of reads from the initial raw libraries. Reference-guided transcriptome based on stringtie merge assembled a total of 276,918 transcripts at 186,948 loci (genes) with an N50 of 1,869. BUSCO assessment against Poales lineage (odb_10) assigned a score of 93.5% (S:12.9%, D:80.6%, F:1.3%, M:5.2%) with 65 and 252 out of 4896 BUSCOs as fragmented and missing respectively, indicating that the assembled transcriptome was of high-quality in terms of completeness. The alignment rate for most of the samples was >72%, with a mean alignment rate of ∼76%. Functional annotation of transcripts based on homology search against local databases of *Triticum aestivum* protein sequences from UniProt, NCBI and ensemble annotated 220,044 (∼79%), 220,817 (∼80%), 215,323 (∼78%) respectively, of the total 276,918 transcripts. The isoform with the highest bitscore at a given locus was used to annotate the gene.

### Differentially expressed genes (DEGs)

#### Differentially expressed genes under ice encasement

A total of 12,142 genes were identified as differentially expressed at different contrasts under long-term ice encasement conditions in timothy; CA vs T1 (2,663 down; 5,100 up), CA vs T2 ( 4,526 down; 4,981 up) and T1 vs T2 (217 down; 49 up). As hypothesized, the DEGs identified under ice encasement stress (Table 2) overlap with some of the core freezing, and hypoxia responses in plants (Bailey-Serres and Voesenek, 2010; Jethva et al., 2022; Pashapu et al., 2024). Some of the DEGs identified include ethylene-responsive transcription factors (ERFs), plant cysteine oxidase (PCOs), prolyl 4-hydroxylase, sucrose non-fermenting-1-related protein kinase (SnRK) (hypoxia signalling), alcohol dehydrogenase (ADH), lactate dehydrogenase (LDH), pyruvate decarboxylase 2 (PDC2), sucrose synthase, pyruvate kinase, ATP-dependent 6- phosphofructokinase (glycolysis), dehydration responsive element-binding 1 (DREB1/C-repeat binding factor (CBF)), late embryogenesis abundant (LEA) proteins, dehydrins, early-responsive to dehydration (ERD) (cold/freezing responses), respiratory burst oxidase homolog protein (RBOH), alternative oxidase (AOX), peroxidases, glutathione S-transferase, superoxide dismutase (SOD) (ROS signalling & homeostasis) (Figure 3, Figure S2, Table 2). A list of all DEGs identified at different contrasts is provided in the supplementary material. GO enrichment analysis revealed that genes upregulated under ice encasement are linked to the terms “response to chemical”, “pyruvate metabolic process”, “response to abiotic stimulus”, “respiratory electron transport chain” (biological processes), “DNA-binding transcription factor activity”, “oxidoreductase activity”, “6-phosphofructokinase activity” (molecular functions) and “cytosol”, “perinuclear region of cytoplasm” (cellular components), while the downregulated genes were linked to the terms “polysaccharide metabolic process”, “carbohydrate metabolic process”, “cell wall organization or biogenesis”, “response to oxidative stress” (biological processes), “peroxidase activity”, “microtubule motor activity”, “transporter activity” (molecular functions) and “extracellular region”, “integral component of membrane”, “cell wall” (cellular components) (Figure S3). Furthermore, KEGG pathway analysis identified that the upregulated DEGs are involved in “Glycolysis / Gluconeogenesis” (taes00010), “Oxidative phosphorylation” (taes00190), “Pyruvate metabolism” (taes00620), “Biosynthesis of amino acids” (taes01230), “carbon metabolism” (taes01200) pathways and the downregulated DEGs are involved in “Phenylpropanoid biosynthesis” (taes00940), “Photosynthesis” (taes00195), “ABC transporters” (taes02010), “Flavonoid biosynthesis” (taes00941) pathways (Figure 4).

**Figure 3:**
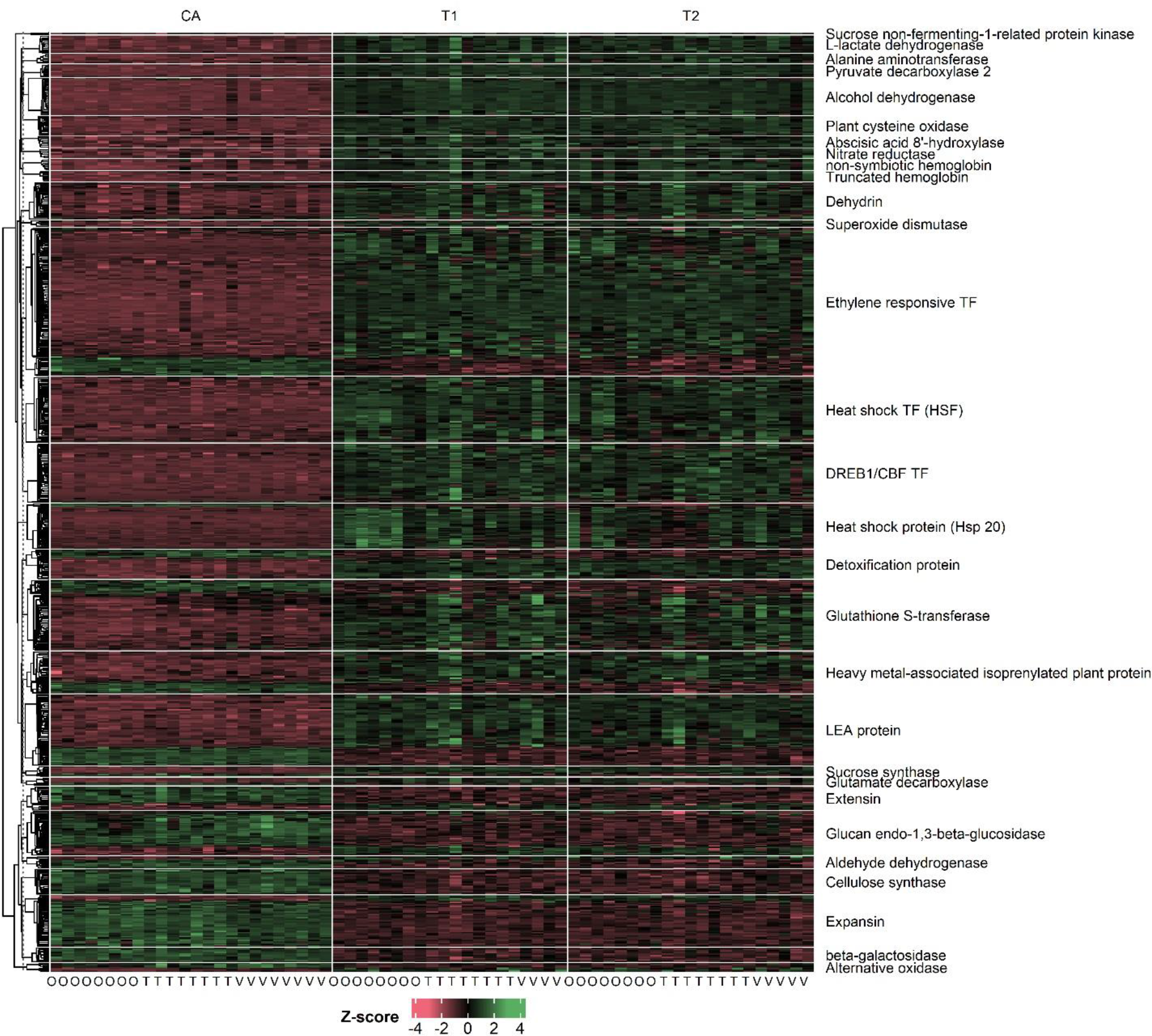
Normalized expression of a few genes identified as differentially expressed under ice encasement. Rows are split based on genes and columns are split based on treatment.

**Figure 4:**
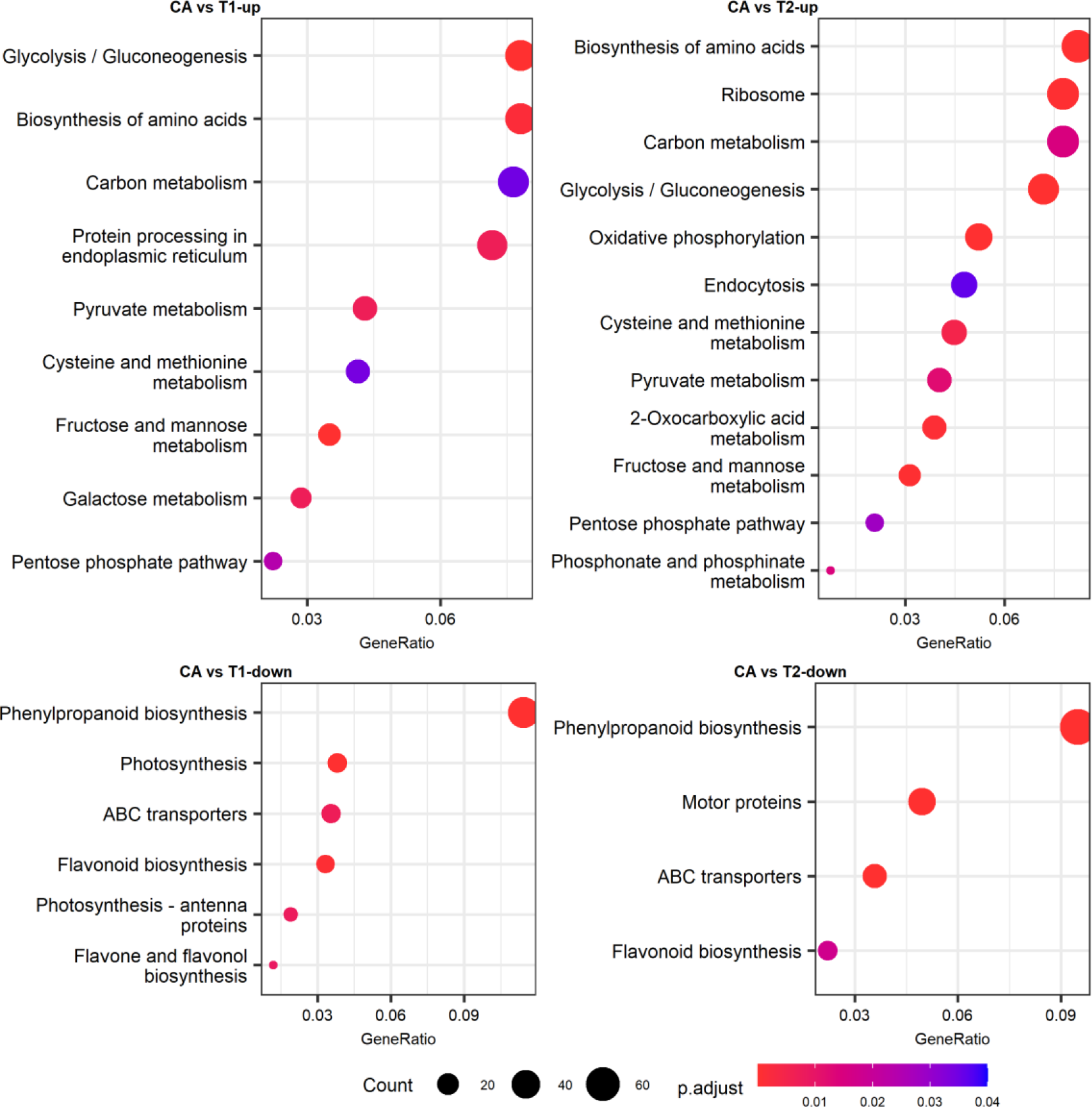
Enriched KEGG pathways under ice encasement in timothy. Up and down denote the differential expression in the contrast.

**Table 2:** A few of DEGs under ice encasement stress in timothy. The list is based on results from current study and literature review on plant stress responses.

| Gene/Protein | Role/Function | Reference | Expression |
| --- | --- | --- | --- |
| 1-aminocyclopropane-1-carboxylate (ACC) synthase | Conversion of S-adenosyl-L-methionine to ACC | (Yang and Hoffman, 1984) | Up |
| 1-aminocyclopropane-1-carboxylate (ACC) oxidase 1 like | Catalyzes the conversion of ACC to ethylene | (Hamilton et al., 1991) | Up |
| Ethylene receptor 2 | Ethylene signaling; overexpression suppresses alpha-amylase | (Wuriyanghan et al., 2009) | Up |
| Ethylene response sensor 2 | Ethylene signalling | (Wuriyanghan et al., 2009; Zhao et al., 2021) | Down |
| Ethylene insensitive 3 | Ethylene signalling | (Chao et al., 1997) | Up |
| Absciscic acid 8'-hydroxylase | Catalyzes oxidation of ABA to phaseic acid (PA) under submergence | (Saika et al., 2006) | Up |
| Plant cysteine oxidase (PCOs) | Degradation of ERF VIIs under normoxia | (White et al., 2017) | Up |
| Ethylene-responsive transcription factors (1, ERF071, ERF073, RAP2-3, RAP2-6) | O2 signalling, regulation of hypoxic stress-responsive genes | (Licausi et al., 2010; Yang et al., 2011) | Up |
| Prolyl 4-hydroxylase | Low oxygen sensing | (Konkina et al., 2021) | Up |
| Mitochondrial ATPase inhibitor | Inhibits mitochondrial ATP synthase activity | (Norling et al., 1990) | Up |
| Sucrose non-fermenting-1-related protein kinase | Energy sensing: positive regulation of anaerobic respiration | (Emanuelle et al., 2016; Luo et al., 2022) | Up |
| Serine/threonine protein kinase TOR (only at CA vs T2) | Regulates hypoxic gene expression based on ATP availability | (Kunkowska et al., 2023) | Down |
| Serine/threonine-protein kinases (SAPK3, SAPK4) | Stress-responsive gene expression | (Diédhiou et al., 2008; Lou et al., 2023) | Up |
| Auxin responsive factor | Regulation of auxin-responsive genes | (Tiwari et al., 2003) | Down |
| Auxin-responsive proteins (SAUR, IAA) | Regulation of development, growth and stress responses | (Ren and Gray, 2015; Stortenbeker and Bemer, 2019) | Up/Down |
| 3-ketoacyl-CoA synthase | Very-long-chain fatty acid synthesis | (Xie et al., 2015) | Up/Down |
| Acyl-CoA dehydrogenase | Fatty acid degradation | (Arent et al., 2008) | Up |
| Phosphatidic acid phosphatase | Lipid metabolism and homeostasis | (Carman and Han, 2019) | Up |
| Ferredoxin-3 chloroplastic-like | Photosynthesis | (Fukuyama, 2004) | Up |
| Chlorophyll a-b binding protein | Photosynthesis; light harvesting | (Liu et al., 2013) | Down |
| Photosystem I | Photosynthesis: light-driven transmembrane electron transfer | (Melkozernov et al., 2006) | Down |
| Chlorophyll b reductase NYC1 | Degrades chlorophyll under high-light darkness | (Sato et al., 2015) | Down |
| Phytochrome-interacting transcription factor 13 | Light signal transduction, role in growth and development | (Zhang et al., 2017) | Up |
| FAR1-related sequence protein | Light signal transduction | (Wang and Wang, 2015) | Down |
| Pseudo-response regulator | Regulates expression of circadian clock genes | (Farré and Liu, 2013) | Up |
| Pyruvate decarboxylase (PDC) 2 | Fermentative pathway: converts pyruvate to acetaldehyde | (Ismond et al., 2003) | Up |
| Alcohol dehydrogenase (ADH) (1 and 3-like) | Fermentative pathway: converts acetaldehyde to ethanol | (Ismond et al., 2003) | Up |
| Lactate dehydrogenase | Fermentative pathway: converts pyruvate to lactate | (Dolferus et al., 2008) | Up |
| Pyruvate kinase | Glycolysis | (Hattori et al., 1995) | Up |
| Glyceraldehyde-3-phosphate dehydrogenase | Glycolysis | (Yang et al., 1993) | Up |
| Glucose-6-phosphate isomerase | Glycolysis; conversion glucose-6-phosphate to fructose-6-phosphate | (Dolferus et al., 1997) | Up |
| Fructose-bisphosphate aldolase | Glycolysis | (Dolferus et al., 1997; Lu et al., 2012) | Up |
| Sucrose synthase | Sugar metabolism | (Stein and Granot, 2019) | Up |
| alpha-amylase | Starch degradation | (Loreti, 2003) | Up |
| beta-amylase (3, and 9-like) | Starch degradation | (Monroe et al., 2014; David et al., 2022) | Up |
| Fructan 6-exohydrolase | Hydrolysis of fructans | (Lothier et al., 2014) | Up |
| beta-fructofuranosidase | Hydrolysis of sucrose to glucose and fructose | (Roitsch and González, 2004) | Up |
| ATP-dependent 6-phosphofructokinase (PFK) | Phosphorylation of fructose-6-phosphate to fructose-1,6-bisphosphate | (Mustroph et al., 2013) | Up |
| Pyrophosphate--fructose 6-phosphate 1-phosphotransferase (PFP) | Phosphorylation of fructose-6-phosphate to fructose-1,6-bisphosphate | (Mustroph et al., 2013) | Up |
| Respiratory burst oxidase homolog protein (A-like, B-like) | ROS signalling | (Suzuki et al., 2011; Yang et al., 2017) | Up |
| Superoxide dismutase | Detoxification of ROS | (Alscher, 2002) | Up |
| Aldehyde dehydrogenase | Detoxification of aldehydes and ROS scavenging | (Kotchoni et al., 2006) | Down |
| Glutathione S-transferase | ROS scavenging | (Vijayakumar et al., 2016) | Up/Down |
| Glutathione peroxidase (only in CA vs T1) | ROS signalling and scavenging | (Bela et al., 2022) | Up |
| Glutathione hydrolase (γ-glutamyl transpeptidase) | Detoxification | (Martin et al., 2007) | Down |
| Glutathione reductase | ROS scavenging | (Gill et al., 2013) | Up |
| Alternative oxidase 1a (1c is downregulated) | Alleviate oxidative stress by regulating ROS levels | (Maxwell et al., 1999; Saha et al., 2016) | Up |
| Catalase | ROS scavenging | (Mhamdi et al., 2010) | Down |
| Plant peroxidases | ROS homeostasis and lignin biosynthesis | (Yoshida et al., 2003; Passardi et al., 2005) | Down |
| Aldo-keto reductase | Detoxification of aldehydes and ketones | (Simpson et al., 2009) | Up |
| Multi-antimicrobial extrusion protein | Detoxification | (Santos et al., 2017) | Up |
| Metal tolerance protein | Metal tolerance and ion homeostasis | (Das et al., 2022) | Up |
| Heavy metal-associated isoprenylated plant protein | Metal detoxification (metallochaperones) & stress response | (khan et al., 2019; Rono et al., 2022) | Up/Down |
| ATP binding cassette (ABC)-transporters | Transport of various substrates & regulation of stress response | (Do et al., 2021) | Up/Down |
| NADH dehydrogenase | Oxidative phosphorylation | (Ramírez-Aguilar et al., 2011) | Up |
| NADH-ubiquinone oxidoreductase | Oxidative phosphorylation | (Subrahmanian et al., 2016) | Up |
| Cytochrome c oxidase | Oxidative phosphorylation, ROS homeostasis | (De Santis et al., 1999; Analin et al., 2020) | Up |
| Nitrate reductase | Reduction of nitrate to nitrite, enhances hypoxic stress tolerance | reviewed by (Timilsina et al., 2020, 2022) | Up |
| Ferredoxin--nitrite reductase | Reduction of nitrite to ammonia | reviewed by (Timilsina et al., 2020) | Up |
| non-symbiotic hemoglobin | Nitric oxide (NO) detoxification | (DORDAS et al., 2003; Hebelstrup et al., 2012) | Up |
| Truncated hemoglobin Hb2b | Might play a role in the NO-dependent pathway under hypoxia | (DORDAS et al., 2003; Hemschemeier et al., 2013) | Up |
| Tubulin alpha chain & beta chain | Constitutes of the cytoskeleton (stress response) | (Ridha Farajalla and Gulick, 2007; Kumar et al., 2022) | Down |
| Kinesin-like proteins | Motor proteins; cell division, development | (Nebenführ and Dixit, 2018) | Down |
| Myosin | Motor proteins | (Nebenführ and Dixit, 2018) | Down |
| Aquaporins (NIP, TIP, PIP) | Transport of various molecules | (Tan et al., 2018) | Down |
| beta-galactosidase | Breakdown of cell wall polysaccharides | (Pandey et al., 2017) | Down |
| Expanin | Cell wall remodelling | (Fukao et al., 2006; Liu et al., 2019) | Down |
| Cellulose synthase | Cellulose biosynthesis | (Speicher et al., 2018) | Down |
| Arabinogalactan proteins (1-like=up, fasciclin-like=down) | Maintaining cell wall integrity and remodelling | (Mareri et al., 2019; Rui and Dinnyen, 2020) | Up/Down |
| Pectinesterase | Cell wall remodeling; de-esterification of pectin | (Wu et al., 2018) | Down |
| Polygalacturonase | Cell wall remodelling; pectin degradation | (Liu et al., 2014) | Down |
| Xyloglucan endotransglucosylase/hydrolase | Cell wall remodeling | (Genovesi et al., 2008; Takahashi et al., 2021) | Up/Down |
| Xylanase inhibitor protein | Cell wall-associated protein involved in defense responses | (Tundo et al., 2022) | Up |
| Growth-regulating factor 3-like | Developmental processes and stress response | (Piya et al., 2020) | Up |
| Cell number regulator (8,10) | Regulate organ size | (Guo et al., 2010) | Up |
| LOB domain-containing protein; 37, 40, 41 | Regulate plant organs and other developmental processes | (Zhang et al., 2020) | Up |
| Serine/threonine-protein kinase TOUSLED | Regulate cell cycle progression through chromatin dynamics | (Carrera et al., 2003) | Up |
| Chaperone protein DnaJ-like | Co-chaperones; protein homeostasis | (Pulido and Leister, 2018) | Up |
| Drought-induced protein (19-like, 1) | ABA-dependent signalling pathway & drought tolerance | (Wu et al., 2022a; Luo et al., 2023) | Up |
| Dehydrin | Stabilize membranes & maintain cellular homeostasis under | (Liu et al., 2017; Yu et al., 2018) | Up |
| Late embryogenesis abundant proteins | Cryoprotectant & maintaining cellular homeostasis under stress | (Hinch and Thalhammer, 2012) | Up/Down |
| DREB1/CBF TF | Regulation of cold stress-responsive genes | (Akhtar et al., 2012; Hwarari et al., 2022) | Up |
| Early-responsive to dehydration 15-like | Regulation of stress-responsive genes; drought, cold | (Alves et al., 2011) | Up |
| Heat shock TF | Regulation of heat shock proteins | (Guo et al., 2016; Kumar et al., 2021) | Up |
| Heat shock proteins (HSPs): 20, 70, 90 | Molecular chaperones; enhance stress responses | (ul Haq et al., 2019) | Up |
| Universal stress protein A (USPA) family | Acts as chaperones under heat, cold and oxidative stress | (Jung et al., 2015; Melencion et al., 2017) | Up |

#### Differential gene expression under ice encasement in field survivors vs. original material

Comparative transcriptomic analysis between field survivors and plants raised from the original seed lots under different ice encasement treatments identified a total of 2,363 differentially expressed genes; CA (171 down and 1,333 up), T1 (244 down and 383 up), T2 (344 down and 383 up). Given that cold-acclimated (CA) and non-ice encased timothy samples were used as controls, the primary focus is on the DEGs at comparisons under ice encasement conditions i.e., at T1 and T2. Genes involved in some of the core freezing and hypoxic stress were observed to be differentially expressed under ice encasement conditions. Ethylene-responsive transcription factors, which are known to be key regulators of gene expression under hypoxic conditions (Licausi et al., 2010), had lower expression in field survivors compared to plants of the original material. In particular, ethylene-responsive transcription factor (ERF) 1-like had 6 and 12-fold lower expression in field survivors at T1 and T2, respectively. Similarly, ERF071-like, ERF073- like and ERF RAP2.13 also had more than a 3-fold lower expression in the field survivors relative to plants from the original material at T2. Dehydrin, which is known to be involved in freezing stress responses had contrasting expression patterns at T1 (8-fold upregulated) and T2 (6-fold downregulated) in field survivors. The expression levels of glyceraldehyde-3-phosphate dehydrogenase, pyruvate kinase isozyme A, and fructose-1,6-bisphosphatase, which are known to be involved in glycolysis, were relatively higher in field survivors at T2. Apart from these, genes coding for glutathione s-transferase, cinnamyl-alcohol dehydrogenase, cytochrome P450, MADS-box TFs, Aldo-keto reductase 3, BURP domain, cysteine-rich receptor-like protein kinase (CRKs), terpene synthase and several genes involved in plant immunity and defence response; pathogenesis-related proteins, Protein WIR1A-like (Bull et al., 1992), glucan endo-1,3-beta-glucosidase (Cheong et al., 2000), DIBOA-glucoside dioxygenase (Shavit et al., 2022), and calmodulin-binding protein 60 (CML60) (Truman et al., 2013) had elevated expression in field survivors compared to the original material (Figure S4). GO enrichment analysis revealed that genes with elevated expression in field survivors are linked to “diterpenoid biosynthetic process”, “response to biotic stimulus”, “polyketide metabolic process” (biological processes), “extracellular region” (cellular components), “glutathione transferase activity”, “oxidoreductase activity”, “hydrolase activity” (molecular functions), while genes with lower expression were linked to “cell wall macromolecule metabolic process”, “alcohol biosynthetic process” (biological processes), “cell wall”, “apoplast” (cellular components), “structural molecule activity”, “xyloglucan:xyloglucosyl transferase activity” (molecular functions) (Figure S5). Furthermore, KEGG enrichment analysis identified monoterpenoid biosynthesis (taes00902), diterpenoid biosynthesis (taes00904), “Glutathione metabolism (taes00480)” and Ribosome (taes03010) pathways as enriched under ice encasement conditions.

#### Differences between northern and southern adapted cultivars under ice encasement

The northern-adapted cultivar Engmo and the southern-adapted cultivar Grindstad displayed varying levels of ice encasement tolerances in the original material and varying levels of loss of ice encasement tolerance in field survivors relative to their counterparts from the original material (Table 1). Therefore, we aimed to identify the differences between Engmo and Grindstad in both the original material and the field survivors (from both locations), using the expression levels of Grindstad as baseline at a given comparison. Comparative transcriptomic analysis identified a total of 1,346 DEGs between Engmo and Grindstad under ice encasement conditions (T1 and T2) across all comparisons of interest. In the original material, we identified 89 and 108 genes as having elevated expressions in Engmo at T1 and T2, respectively, while 35 and 129 genes had lower expressions at T1 and T2, respectively. Meanwhile, in field survivors, we identified 266 and 395 genes as upregulated in Engmo at T1 and T2, respectively, while 193 and 352 genes were downregulated at T1 and T2, respectively. In the original material, glutamate decarboxylase, superoxide dismutase, heat shock protein (HSP) 70 and 90, glutathione s-transferase, glyceraldehyde-3-phosphate dehydrogenase and dehydrin had elevated expression at T2 in Engmo, while the expression of histone 2, histone 3, cinnamyl-alcohol dehydrogenase, plant peroxidase, nitrate reductase, heavy metal-associated isoprenylated plant protein and serine/threonine-protein kinase STY13-like was higher in Grindstad at T2 (Figure S6A). In field survivors, the expression of dehydrin, LEA protein, low-temperature-induced protein, NAD(P)H-quinone oxidoreductase, phosphoenolpyruvate carboxylase, cinnamyl-alcohol dehydrogenase, xyloglucan endotransglucosylase/hydrolase, glyceraldehyde-3-phosphate dehydrogenase, 4- hydroxyphenylpyruvate dioxygenase, phosphoenolpyruvate carboxylase, enoyl-CoA reductase and NAD kinase was elevated in Engmo. Interestingly, genes encoding 3-ketoacyl-CoA synthase, very-long-chain 3-oxoacyl-CoA reductase, fatty acyl-CoA reductase, chlorophyll a-b binding protein, photosystem II, oxygen-evolving enhancer protein, sucrose synthase, histone 2 and histone 3 seem to elevated expression at T2 relative to T1 in Engmo, while their expression largely remained unchanged in Grindstad (Figure S6B).

## Discussion

### Ice encasement stress responses in plants

Restriction of gaseous exchange between the plant and the surrounding environment due to the formation of ice cover and the subsequent emergence of hypoxic conditions marks the onset of ice encasement stress. Therefore, like under flooding/submergence (Licausi et al., 2011; Sasidharan and Mustroph, 2011), sensing low oxygen is likely the first step by which overwintering plants initiate ice-encasement stress responses. It is important to remember that ice encasement events usually occur mid-winter when overwintering plant species must have already undergone metabolic readjustments to contend with freezing stress. The reprogramming of the cellular metabolism under hypoxic conditions is primarily driven by the group VII ethylene-responsive factor (ERF) transcription factors, whose stabilization under low oxygen conditions leads to transcriptional activation of core anaerobic response genes (ARGs) (Licausi et al., 2011; Sasidharan and Mustroph, 2011), which include alcohol dehydrogenase (ADH), pyruvate dehydrogenase (PDC), lactate dehydrogenase (LDH) and sucrose synthase (SUS). These ARGs avoid the over-reduction of the NAD^+^/NADH pool and help in addressing severe limitations in ATP synthesis by mitochondrial respiration through glycolytic fermentation (Perata and Alpi, 1993; Lee et al., 2011; Jethva et al., 2022). Similar to submergence, expression of ERF TFs, ADH, PDC, LDH, alanine aminotransferase (AAT) and SUS was highly elevated under ice encasement conditions (Table 2 and Figure 3). Due to the lack of sunlight (at higher latitudes) and partial or complete shutdown of the photosynthetic apparatus (Boese and Huner, 1990; Strand et al., 1997; Hajihashemi et al., 2018) during winters, the metabolic processes rely on the carbohydrate reserves. In line, genes involved in carbohydrate and sugar metabolism; SUS, glyceraldehyde-3- phosphate dehydrogenase (GAPDH), pyruvate kinase (PK), pyruvate phosphate dikinase (PPDK), alpha-amylase, beta-amylase, glucose-6-phosphate isomerase, and hexokinase were highly upregulated under ice encasement conditions. The observation that the expression of these genes remains elevated even at T2 (long-duration ice encasement) supports the claim that carbohydrate depletion is not the main reason for plant mortality under ice encasement (McKersie et al., 1982). Unlike under submergence, plant death due to energy starvation or carbohydrate depletion is highly unlikely under ice encasement. This is because i) overwintering plants accumulate carbohydrate reserves during cold acclimation (Kaplan et al., 2004) to survive the long winters; and ii) lower metabolic and respiration rates (Ögren, 2000; Gudleifsson, 2009) due to freezing conditions drastically reduce the consumption of carbohydrate reserves.

Unlike under submergence, encasement in ice prevents the continuous leaching of the anaerobic byproducts and previous studies have shown that accumulation of these byproducts to toxic levels is the primary reason for plant death under ice encasement (Andrews and Pomeroy, 1979; Andrews, 1997). The formation and accumulation of anaerobic byproducts is a function of the anaerobic respiration rate. Therefore, ice-encased plants could prevent the accumulation of anaerobic byproducts to toxic levels by i) imposing strong metabolic quiescence; ii) employing alternative processes that produce anaerobic byproducts in lower quantities for ATP synthesis; and iii) detoxification. Studies by Andrews (1997), Bertrand et al. (2001) and Höglind et al. (2010) revealed that timothy accumulated less anaerobic byproducts while maintaining relatively high water-soluble carbohydrate (WSC) content, indicating that the superior ICET of timothy is due to its slower glycolytic metabolism compared to other grass species. Maintaining low glycolytic rates under ice encasement requires tight regulation of essential metabolic processes (metabolic quiescence), and how species with superior ICET are able to maintain low glycolytic rates is not yet understood. Weits et al., 2014 showed that in *Arabidopsis thaliana*, plant cysteine oxidase 1 (PCO) and PCO2 are hypoxia-inducible and redundantly repress the anaerobic gene expression while themselves being the targets of ERF-VII transcription factors. In the current study, plant cysteine oxidase 2 and 3 (PCO), were upregulated (2 – 12-fold) in comparisons CA vs T1 and CA vs T2 (Figure 3), this observation indicates that anaerobic responsive gene expression in timothy is tightly regulated. Plant mitochondria can preserve their structure and function even under anoxic conditions, in particular when exposed to nitrate (Fox and Kennedy, 1991; Vartapetian, 2003). Studies on mitochondria isolated from roots of barley, rice and pea seedlings demonstrated that plant mitochondria have the capacity to use nitrite as an electron acceptor under oxygen-deficient conditions to drive ATP synthesis, and this also led to decreased production of reactive oxygen species (ROS) and decreased lipid peroxidation (Igamberdiev, 2004; Stoimenova et al., 2007; Gupta et al., 2017). The expression of genes involved in nitrite-driven anaerobic ATP synthesis i.e., nitrate reductase, ferredoxin-nitrite reductase, non-symbiotic hemoglobin, and truncated hemoglobin were upregulated in timothy at T1 and remained stable at T2 (Figure 3). Only the expression of non-symbiotic hemoglobin, involved in NO scavenging (Hebelstrup et al., 2012), was observed to increase from T1 to T2 (not significantly DE). It would be interesting to investigate the extent to which ATP needed to fuel the cellular processes under ice encasement conditions is produced through nitrite-driven anaerobic ATP synthesis in high ICET and low ICET species. In contrast to the previous strategies focusing on reducing the production of anaerobic byproducts, the last strategy involves adeptly scavenging the anaerobic byproducts. Under ice encasement plants mainly accumulate CO_2_, ethanol, lactate, and malate, pyruvate, citrate and fumarate in smaller amounts (Gudleifsson, 1994, 2010; Bertrand et al., 2001). A study by Andrews and Pomeroy (1979) in winter wheat demonstrated that CO2, ethanol and a combination of these agents in particular are the major contributors to reduced plant viability under ice encasement with cell membrane being the site of damage. Furthermore, oxygen deprivation disrupts a host of cellular processes resulting in cytosolic acidification as well as accumulation of ROS and reactive nitrogen species (RNS) that potentially lead to cell damage (Hebelstrup and Møller, 2015; Turkan, 2018; Jethva et al., 2022). Therefore, regulating the levels of ROS and RNS and preventing the accumulation of anaerobic byproducts to toxic levels are crucial to avoid irreversible damage. In the current study, the expression of genes involved in detoxification; superoxide dismutase (SOD), glutathione S-transferase (GST), glutathione reductase (GR), alternative oxidase (AOX) 1a, ADH, aldo-keto reductase (AKR), metal tolerance protein, heavy metal-associated isoprenylated protein, multi-antimicrobial extrusion protein was upregulated at CA vs T1 and CA vs T2 (refer Table 2). Interestingly, the expression of a few detoxification genes; aldehyde dehydrogenase, AOX 1c, glutathione hydrolase, catalase (CAT), and plant peroxidase, short-chain dehydrogenases (SDR) was downregulated under ice encasement. In view, of these observations, the tight regulation of anaerobic gene expression by PCOs, the utilization of nitrite-driven anaerobic ATP synthesis and the selective regulation of genes involved in detoxification might be the mechanisms contributing to prolonged survival of timothy under ice encasement.

Plant species frequently experiencing flooding or submergence employ two different strategies to protect against damage: i) low oxygen escape strategy (LOES) and ii) low oxygen quiescence strategy (LOQS). In LOES plants rapidly elongate stems, petioles or leaves to restore contact with air, while LOQS involves readjusting the metabolism (Bailey-Serres and Voesenek, 2008). It is highly unlikely that plants employ LOES under ice encasement conditions as it necessitates initiating growth under freezing conditions, implying deacclimation, which is rather detrimental to plant survival. Additionally, unlike water the ice surface is impervious, therefore, readjusting the metabolism to maintain metabolic quiescence (LOQS) is more likely to be the main strategy to cope with the stress induced by ice encasement. Upregulation of genes controlling cell cycle and organ size, i.e., growth-regulating factor 3-like, cell number regulator, lateral organ boundary (LOB) domain-containing protein, and TOUSLED serine/threonine-protein kinase, and downregulation of many genes linked to cell wall modification and growth, i.e., expansins, cellulose synthase, xyloglucan endotransglucosylase/hydrolase, fasciclin-like arabinogalactan proteins, pectin esterase, and polygalacturonase support a LOQS response under ice encasement (Table 2). Moreover, the expression of DREB1/CBF TFs, dehydrins, LEA proteins, early-responsive to dehydration 15-like, heat shock TFs and heat shock proteins, core genes involved in freezing stress responses of timothy (Pashapu et al., 2024) were observed to be upregulated under ice encasement (Table 2), supporting our initial hypothesis. It would be interesting to compare the expression of these genes under freezing and ice encasement stress.

### Gene expression differentiation between field survivors and original material

Investigating the factors leading to genetic differentiation between plant populations subjected to natural selection in field conditions and plants raised from the seed lots used to establish those field trials, is key for understanding and predicting species response to changing climatic conditions and developing new plant varieties with better persistence. The prevalence and the intensity of a given stress, the standing genetic variation of the population and the temporal duration collectively determine the extent of genetic differentiation. Winter survival is a complex trait and plants in the field must cope with multiple winter stresses, often at the same time, thus it is highly extremely difficult to determine the main factor driving selection and the cause of the plant death under field conditions. Previously freezing tolerance was considered as the primary determinant of winter survival (Pulli et al., 1996; Rognli, 2013), but with the predicted increase in WWEs at northern latitudes (Vikhamar-Schuler et al., 2016), tolerance to ice encasement stress might be equally important. Therefore, one of the objectives of the current study was to discern variation in gene expression between field trial survivors and plants raised from seed lots used to establish the field trials under ice encasement stress.

There was a gradual decline in the density of timothy plants in the field plots from the first ley year (2017) to the last (2020) with reductions ranging from about half to two-thirds in most plots, indicating substantial selection pressure. Unfortunately, the prevalence of ice on the field plots was not recorded during the field experiments, therefore the findings of this study cannot be entirely attributed to selection due to ice encasement under field conditions. The stable overexpression group VII ERF TFs is known to positively regulate hypoxic gene expression, thereby enhancing waterlogging tolerance (Gasch et al., 2016; Fukao et al., 2019). The lower ICET (LD_50_) of the field survivors in the current study can be partly explained by the relatively lower expression of ERF 1-like, ERF071-like, ERF073-like and ERF RAP2.13 in field survivors compared to plants from the original material. Moreover, the expression of glyceraldehyde-3- phosphate dehydrogenase, pyruvate kinase isozyme A, and fructose-1,6-bisphosphatase linked to glycolysis (Yang et al., 1993; Hattori et al., 1995) were relatively higher in field survivors at T2. This indicates that the elevated glycolytic rate in field survivors compared to the original material may have depleted carbohydrate reserves while also leading to increased accumulation of anaerobic byproducts. Previous studies in *Arabidopsis thaliana* and *Cistus clusii* showed that older plants and leaves accumulated more H_2_O_2_ and malondialdehyde (Munné-Bosch and Alegre, 2002; Xiang and Rathinasabapathi, 2022) indicating higher oxidative stress in ageing plants and tissues. Thus, the elevated expression of glutathione s-transferase, catalase, aldehyde oxidase, aldo-keto reductase, short-chain dehydrogenase/reductase linked to ROS scavenging and detoxification (Simpson et al., 2009; Mhamdi et al., 2010; Vijayakumar et al., 2016; Wu et al., 2022b) in field survivors compared to the original material might be due to a combination of selection in the field and the age effect. The huge differences in gene expression during cold acclimation (CA), which is crucial for winter survival, could also have led to differences in ICET in field survivors and the original material. We speculate that the lower ICET (LD_50_) of field survivors might be due to differences in CA responses, lower expression of ERF TFs, higher glycolytic rate, and elevated oxidative stress. Further studies should be aimed at investigating the effect of plant age on stress responses in perennial grasses.

### Differences between northern and southern adapted cultivars in field survivors and original material

Persistence for multiple years and higher tolerance to various winter stresses (a prerequisite for persistence) are highly desirable traits in overwintering perennial grasses since the swards need to be re-established less frequently leading to lower production costs. Thus, in the context of perennial grasses, a cultivar with high and relatively stable stress tolerance for multiple years (throughout plant life) is the ultimate breeding goal of plant breeders. In the current study, the northern adapted cultivar Engmo was observed to be more ICET in both the original material and the field survivors, while the southern adapted cultivar Grindstad displayed remarkably stable, albeit lower ICET in both the original material and the field survivors (Table 1). Therefore, we aimed at finding differences in gene expression in Engmo and Grindstad in the original material and field survivors, which might explain the different characteristics of these cultivars.

The higher ICET of Engmo compared to Grindstad observed in the current study is in accordance with the findings of Höglind et al., 2010. Based on the number of DEGs, there were more differences between Engmo and Grindstad in the field survivors than in the original material. In the original materials, contrasting expression of several ice-encasement-responsive genes between the cultivars was observed. The expression of glutamate decarboxylase, superoxide dismutase, heat shock protein (HSP) 70 and 90, glutathione s-transferase, glyceraldehyde-3- phosphate dehydrogenase and dehydrin increased from T1 to T2 in Engmo while their expression decreased in Grindstad (Figure S6A). This observation could partly explain the higher ICET of Engmo in the original material. Results from Höglind et al., 2010 suggested that the higher ICET of Engmo is due to a slower metabolic rate indicated by a slower decline of WSC content compared to Grindstad. As in the original material, expression of several stress-responsive genes was higher in field survivors of Engmo compared to Grindstad, which could explain its higher ICET tolerance (Figure S6B). However, results of the current study cannot explain the decline in ICET in Engmo and the relatively stable expression of ICET in Grindstad among field survivors when compared to their corresponding original material. A similar trend was observed in one of our previous studies investigating freezing stress responses in the same plant material (Pashapu et al., 2024). The differences in their stress tolerances and the contrasting ability to maintain stress tolerances throughout different stages of plant development make Engmo and Grindstad the ideal candidates to study the effect of plant age on stress responses.

## Conclusions

The results of the current study indicate ice encasement stress responses in perennial grasses are fine-tuned to cope with multiple stressful conditions under ice cover and therefore complex. The findings highlighted the presence of a substantial overlap of ice encasement stress responses with those of hypoxic and freezing stresses supporting our initial hypothesis. The stable expression of many genes involved in carbohydrate and sugar metabolism even at T2 (relative to T1) supports the observation of McKersie et al., 1982, that depletion of carbohydrate reserves is not the main reason for plant mortality under ice encasement. The tight regulation of gene expression through induction of PCOs, using nitrite-driven anaerobic pathway for ATP synthesis and selective upregulation of detoxification genes indicate the capacity for flexible metabolic readjustment of timothy to cope with oxygen and energy shortages. Future studies should compare the gene expression of species with different ice encasement tolerances to gain further insights into the mechanisms leading to higher ICET tolerance of timothy. In addition to studying gene expression future studies should also consider measuring the WSC, fructan content, anaerobic metabolite content and changes in gene expression during reoxygenation; which is also considered a major stressor leading to plant mortality.

## Author contributions

S.D, M.J, O.A.R and M.R.K were involved in planning the experiment. S.D and M.J carried out the freezing tests. A.R.P performed bioinformatic data analysis and wrote the manuscript. All co-authors provided critical feedback and helped to finalize the manuscript.

## Acknowledgements

This study was funded by the Norwegian Research Council, project number 303258, “NexTim – Securing adaptation of timothy cultivars under climate change and during seed multiplication using genomics and big-data approaches” and a PhD scholarship to A.R.P from the Faculty of Biosciences (BIOVIT), Norwegian University of Life Sciences (NMBU). We would like to thank Maxime Clausses for RNA extractions and Justine Lèa Marie Leleu for RNA extractions and analyzing part of the data for her master’s thesis.

## Data availability

The RNA-seq data generated is available at EMBL-EBI with accession number E-MTAB-13705

